# FunSpace: A functional and spatial analytic approach to cell imaging data using entropy measures

**DOI:** 10.1101/2022.06.17.496475

**Authors:** Thao Vu, Souvik Seal, Julia Wrobel, Tusharkanti Ghosh, Mansooreh Ahmadian, Debashis Ghosh

## Abstract

Spatial heterogeneity in the tumor microenvironment (TME) plays a critical role in gaining insights into tumor development and progression. Conventional metrics typically capture the spatial differential between TME cellular architectures by either exploring the cell distributions in a pairwise fashion or aggregating the heterogeneity across multiple cell distributions without considering for the spatial contribution. As such, none of the existing approaches has fully accounted for the heterogeneity caused by both cellular diversity and spatial configurations of multiple cell categories. In this article, we propose an approach to leverage the spatial entropy measures at multiple distance ranges to account for the spatial heterogeneity across different cellular architectures. Then, functional principal component analysis (FPCA) targeting sparse data is applied to estimate FPC scores which are then predictors in a Cox regression model to investigate the impact of spatial heterogeneity in the TME on survival outcome, holding other clinical variables constant. Using an ovarian cancer dataset (n = 114) as a case study, we found that the spatial heterogeneity in the TME immune compositions of CD19+ B cells, CD4+ T cells, CD8+ T cells, and CD68+ macrophages, had a significant non-zero effect on the overall survival (p = 0.027). In the simulations studies under different spatial configurations, the proposed method demonstrated a high predictive power by accounting for both clinical effect and the impact of spatial heterogeneity.

## Introduction

The emergence of tumor microenvironment (TME) studies has revealed a critical role of spatial heterogeneity for gaining insights into tumor initiation, development, progression, invasion, metastasis, and response to therapies (1, 2, 3, 4). The TME is known to be complex and heterogeneous due to continuous cellular and molecular adaptions in the primary tumor and its surroundings, which then allow for tumor growth and proliferation. Increasing evidence suggests that in addition to quantities and types, the spatial architectures of cells within the TME influences survival and response to treatment therapy in numerous cancer types (1). For instance, Wang et al. (3) discovered a high level of heterogeneity in the TME of patients with human lung adenocarcinoma (LUAD) by comparing different tumor sites. Particularly, the authors identified the immunological differences in cell subpopulations between the core, middle, and edge of tumors, such that CD4+ naive T cells located at the core of the tumor had higher activation levels in angiogenesis and expressed more immune checkpoint molecules than those at the tumor edge. In another study about human papillomavirus (HPV)-negative head and neck squamous cell carcinoma (HNSCC) tumors, Blise et al. (1) demonstrated that neoplastic tumor-immune cell spatial compartmentalization, rather than mixing, was associated with longer progression free survival (PFS).

Great progress has been made in studying spatial architecture of cells in the TME owing to advances in single-cell multiplex imaging modalities which provide simultaneous quantification and visualization of individual cells in tissue sections (5, 6, 7, 8, 9, 10). More specifically, multiplex tissue imaging (MTI) (11) methods such as cyclic immunoflourescence (CyCIF) (12), CO-Dectection by indEXing (CODEX) (13), multiplex immunohistochemistry (mIHC) (10, 14), imaging mass cytometry (IMC) (15), and multiplex ion beam imaging (MIBI) (16) are capable of measuring the expression of tens of markers at single-cell resolution while preserving the spatial distribution of cells. As an example, multiplex immunohistochemistry (mIHC) detects and visualizes specific antigens in cells of a tissue section by utilizing antibody-antigen reactions coupled to a flourescent dye or an enzyme (17, 18). Another instance includes multiplexed ion beam imaging (MIBI) (16), which utilizes secondary ion mass spectrometry to image metal-conjugated antibodies. As a result, MIBI enables single-cell analysis of up to 100 parameters without spectral overlap between channels. Altogether, imaging provides an additional dimension of spatial resolution to the single cell signature profiles, which in turn allows researchers not only to study cellular composition but also to make inferences about specific cell-cell interactions.

Metrics that quantify the spatial differences between TME cellular architectures can range from simple density ratios of immune cells to tumor cells within specific tumor regions (e.g., tumor center vs. invasive margin) such as Immunoscore (19), to more complex measures utilizing spatial proximity of specific cell types relative to others in the TME such as mixing scores (8) and cellular neighborhood measures (9). Alternatively, the Ripley’s K-function and its variants could also be employed to characterize any single - cell spatial patterns deviated from the complete randomness at any given distance. However, for a given point pattern of multiple cell types, such spatial summary functions typically operate in a pairwise fashion, which involves all possible comparisons of one cell type to another, but not all types simultaneously.

Shannon entropy (20) initially proposed in Information Theory to measure the heterogeneity in observations, has gained popularity in a wide range of applied sciences such as ecology and geography (21, 22, 23), evolutionary biology (24), landscapes (21, 25), and recently in cancer research (26, 27). For instance, Heindl et al. (26) investigated the microenvironmental diversity of ovary tumors and the corresponding local metastasis sites including omentum, peritoneum, lymph node, and appendix by accounting for the collective characteristics of all cell types simultaneously via Shannon diversity index. Particularly, based on the cell type frequency distribution, a high Shannon score indicates similar proportion of each type while a low value suggests dominance of one cell type. In a different manner, (27) associated the diversity of cell compositions in the tumor ecosystem with patients overall survival on a more granular level. Specifically, after cell phenotyping and classification, the authors divided each image into smaller regions such that Shannon entropy can be calculated for each region based on the frequency distribution of each cell type. Accordingly, a high value of the entropy indicate a heterogeneous environment while the reverse holds for a low entropy value. The distribution of Shannon diversity scores was then used as input for Gaussian mixture model to determine the number of clusters, which was referred to as an ecosystem diversity index (EDI).

While the direct application of Shannon entropy in (26) showed some promising results in ovarian cancer, it does not consider the spatial distribution of cell types. In other words, regardless of how cells of different types are distributed on a given image, the Shannon entropy is the same as long as the proportion of each cell type stays the same. The EDI approach, on the other hand, tries to overcome such challenge by considering small neighborhoods of cells through image tessellation. However, this approach relies heavily on how each image is tessellated (e.g., shape and size) and the chosen number of clusters from fitting the model to obtain the EDI score. Herein, by considering a collection of cells on each image as a marked point pattern, we propose an approach that leverages spatial entropy measures (28) to account for spatial heterogeneity across subjects at certain distance ranges and how such variability impacts a clinical outcome of interest. More precisely, if cells of different types are randomly scattered on an image, the spatial entropy at any given distance would be around zero. On the other hand, the spatial entropy would deviate from zero if spatial patterns of cell types in a given local neighborhood are different from the global pattern. The number of distance ranges is rather limited to ensure entropy can be reasonably calculated for each range. As a result, we utilize functional principal component analysis (FPCA) targeting sparse data to obtain subject-specific FPC scores which capture the spatial heterogeneity of the TME compositions. The FPC scores are then served as predictors in a Cox regression model to investigate the association between spatial heterogeneity in the TME compositions with survival outcome. Using the ovarian cancer dataset (n = 114) as a case study, we found that the spatial heterogeneity in the TME immune compositions of CD19+ B cells, CD4+ T cells, CD8+ T cells, and CD68+ macrophages, had a significant non-zero effect on the overall survival (p = 0.027). In the simulations studies under different spatial configurations, the proposed method demonstrated a high predictive power by accounting for both clinical effect and the impact of spatial heterogeneity.

## Real application results

### Ovarian cancer data

Motivated by recent studies on the immune responses in the ovarian TME (29, 30), we explored the spatial heterogeneity in various immune cell subsets such as T and B cells, tumor-associated macrophages. Fig. 1 (A) illustrates distribution of four different immune cell types including CD19+ B cells, CD4+ T cells, CD8+ T cells, and CD68+ macrophages, in four representative individuals. Utilizing the entropy measures introduced in Section for these four categories, i.e. *I* = 4, the heterogeneity in spatial distributions of these cell types was captured for each individual image. Fig. 1 (B) shows such spatial entropy measures for all 114 subjects in the dataset as a function of inter-cell distances. In particular, there was a high level of variation in spatial entropy values at distances less than 250 *µ*m at which some individuals expressed high entropy values while others had values close to zero.

**Fig. 1.**
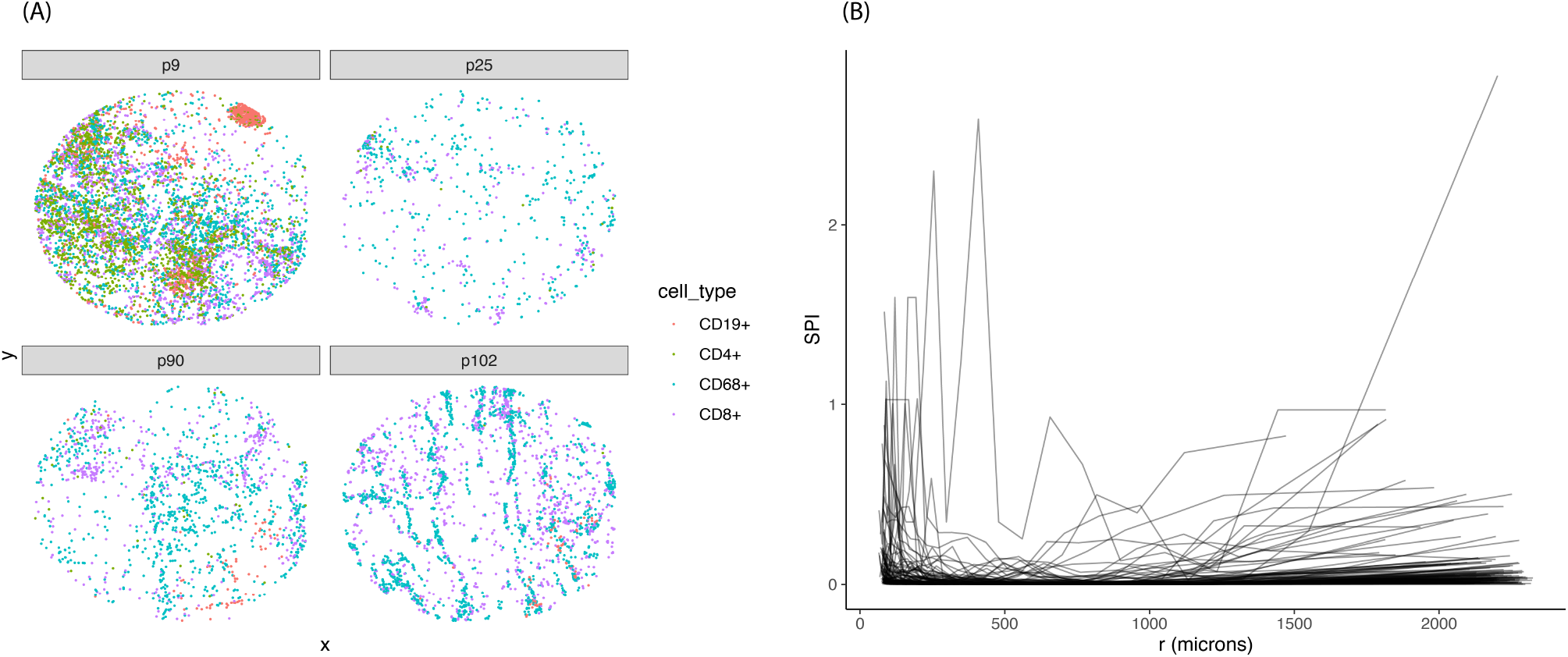
Ovarian cancer dataset: (A) Representative images with distribution of immune cells including CD19+ B cells, CD4+ T cells, CD8+ T cells, and CD68+ macrophages. (B) Spatial entropy of the four immune cell types as a function of inter-cell distances.

**Fig. 2.**
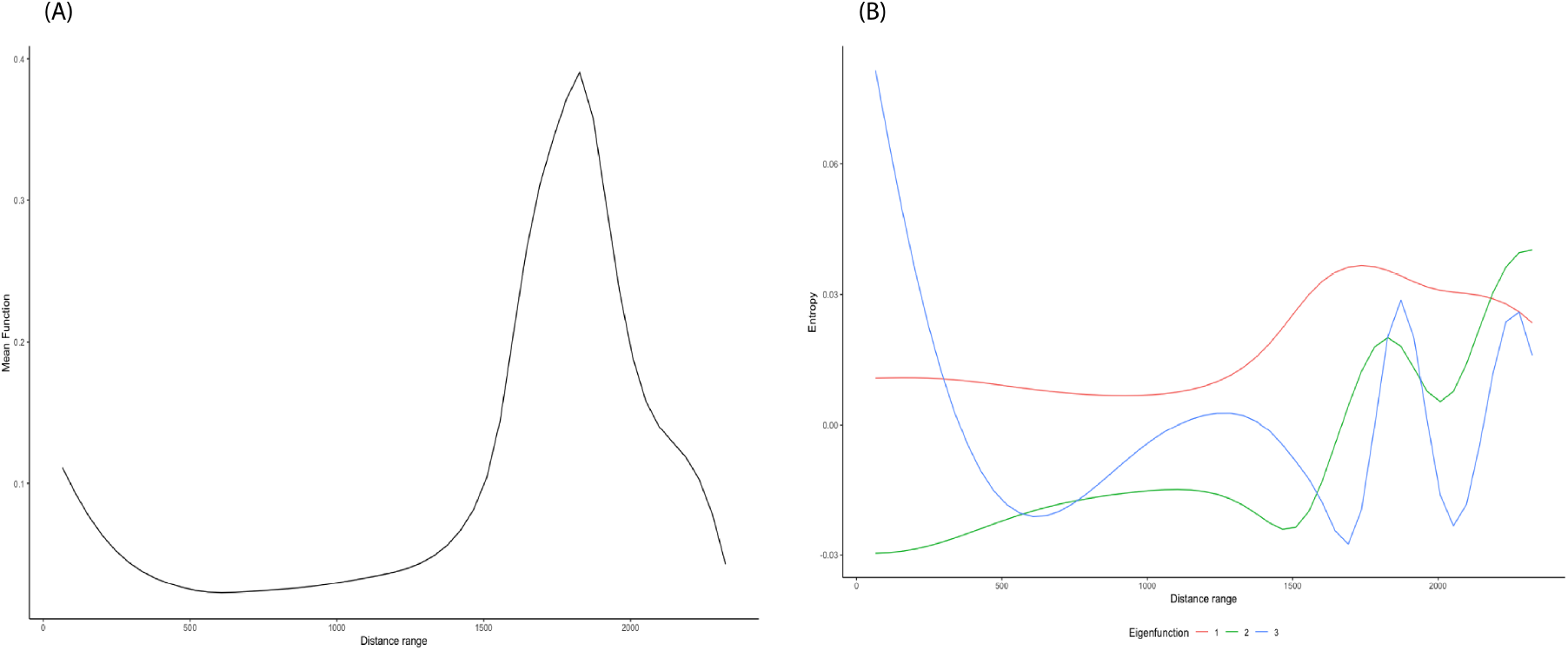
FPCA results from SPI curves in ovarian dataset. (A) Mean function. (B) First three eigenfunctions.

The resulting SPI curves were used as input for the FPC analysis to obtained the estimated FPC scores. Fig. S1 shows the estimated mean function (A) and the first three eigenfunctions (B). The estimated mean function reflected the overall trend starting at relatively high entropy values at short distances (≤ 250 *µm*), then dropping off close to zero for intermediate distances (around 300 − 1000 *µm*). Towards the end of the distance range, the spatial entropy values became larger. The first eigenfunction showed similar trend to the mean function while the second one expressed a contrast in spatial entropy values between the shortest and farthest distances. The third eigenfunction, on the other hand, illustrated a clearer contrast in entropy values between distances of *<* 250 *µm* vs. *>* 250 *µm*. The first three eigenfunctions accounted for 95.6% of the total variation.

By fitting the three selected FPC scores directly into model [4], we investigated the relationship between spatial heterogeneity in distributions of immune cells in the TME and survival outcome, in addition to subject age. Full and restricted models were fit to test the hypothesis [6]. We obtained the p-value of 0.027, indicating the significant nonzero effect of spatial heterogeneity in TME immune compositions on the overall survival.

### TNBC

In this analysis, we focused on exploring the difference in the spatial distributions of endothelial, immune, mesenchymal-like, and tumor cells in the TME. Fig. S2 (A) displays such spatial distributions in four representative images. With the number of categories *I* = 4, the SPI curves were computed for 38 subjects as shown in Fig. S2 (B). For short inter-cell distances (≤200 *µm*), the spatial entropy values were relatively high compared to those between 300 *µm* and 600 *µm*. Specifically, high values of spatial entropy at short distances were due to some individuals having small clusters of immune and tumor cells. As the distances increased, cells of different types started scattering more evenly, leading to the spatial entropy dropping close to zero. Similar to the previous case, we used FPC analysis on the estimated SPI curves to obtain the corresponding FPC scores. Panels (A) and (B) of Fig. S3 display the estimated mean function and first two eigenfunctions, respectively. An overall trend of high entropy values at short distances then dropping off close to zero was reflected in the mean function and the first eigenfunction. In addition, the second eigenfunction depicted a contrast in the spatial entropy values between short distances (≤ 300 *µm*) and distances beyond 600 *µm*. Utilizing model [4], we also tested the relationship between spatial heterogeneity in the distributions of the four cell types in the TME of breast tumor samples and mortality risk. In particular, we fit a full model with the first two FPC scores as predictors in addition to age. A restricted model, on the other hand, only included age as predictor. With limited sample size (*n* = 38), the LRT of the two models resulted in a p-value of 0.31 indicating no significant association.

## Simulation studies

### Setup

We performed simulations studies to evaluate the performance of the proposed approach. We considered a dataset of 100 images (*N* = 100). For simplicity, we considered a total of five cell types (e.g., CD14+, CD19+, CD4+, CD8+, and CK+) per image (i.e., *I* = 5). Total number of cells per type was equally fixed at 600, leading to the total cells *n*_*c*_ = 3000 per image. For each dataset, we assumed that there were two groups of subjects; number of subjects per group followed a binomial distribution with a probability *prob* = 0.5, i.e. *N*_*g*_ ∼ *Binom*(*N, prob* = 0.5) for *g* = 1, 2. We considered two spatial configurations: clustered vs. random (top row of Fig. 3), which subjects in groups 1 and 2 were generated from, respectively. In other words, the proportion of each cell type stayed consistent across subjects, only their spatial distributions varied. The bottom row of Fig. 3 shows the two reference spatial entropy curves corresponding to each configuration. If cells of each type were randomly scattered in a given image, the corresponding spatial entropy values were approximately zero across all distance ranges. Conversely, any configuration deviated from the complete randomness would result in spatial entropy curve above zero.

**Fig. 3.**
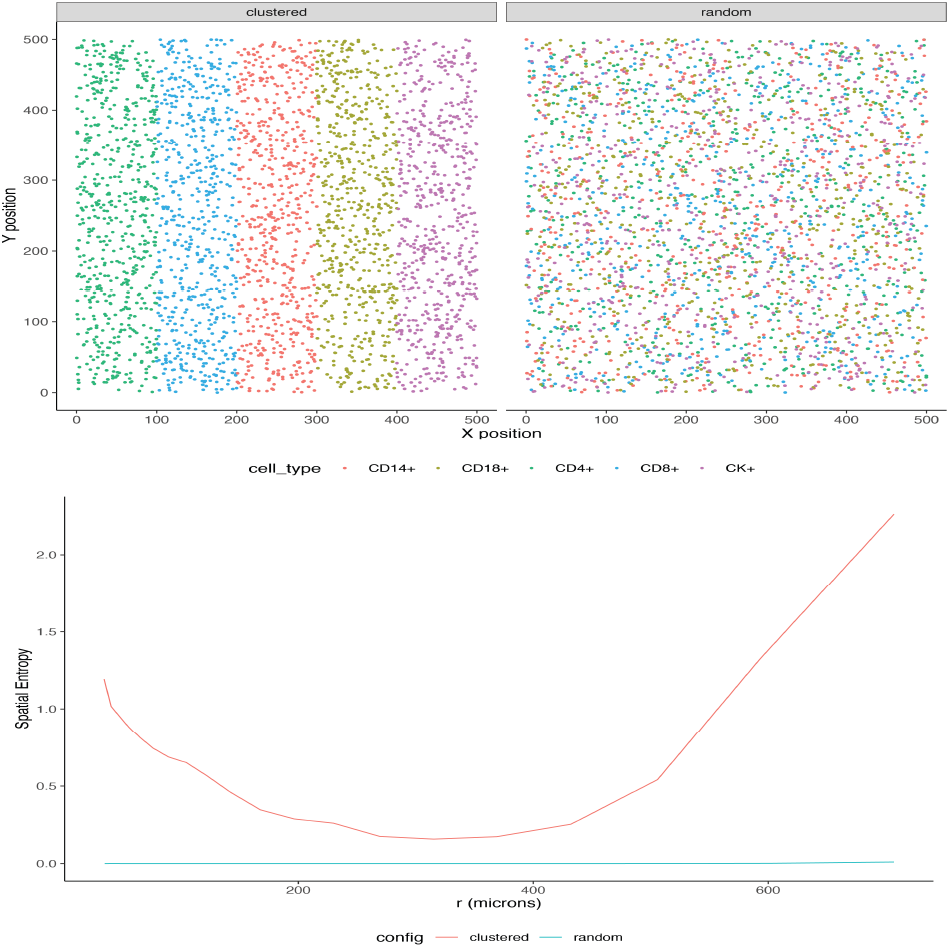
Simulated spatial configurations. Top row: Two reference spatial configurations: clustered (left) and random (right) of five different cell types: CD14+, CD19+, CD4+, CD8+, and CK+. Bottom row: Corresponding spatial entropy at multiple distance ranges for each configuration.

Next, subject-specific entropy curves were generated by adding noise to the reference curves. We constructed three scenarios corresponding to the three levels of additive noise: small, medium, and large (Fig. S4, Supplementary Information). FPCA was performed on the simulated spatial entropy curves to obtain the estimated FPC scores 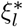 (Section). Again, the number of FPCs *L*^∗^ was chosen such that at least 95% of the total variation was accounted for. Additionally, the scalar covariate 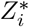 was simulated from the normal distribution with mean and standard deviation obtained empirically from the distribution of age. After mean centering and unit scaling 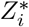 and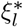, the linear predictor was simulated as 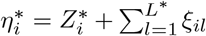 From the fitted model in the real data analysis, we obtained the estimated cumulative baseline hazard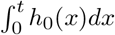, which was then used to generate a survival function for each individual, such that 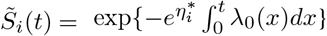. The estimated survival times 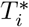 were generated from the survival function; and the censoring times 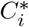 were simulated based on the empirical distribution of the observed censoring times.

### Predicted Performance

At each level additive noise, four datasets of different sizes (N = 100, 200, 500, and 1000, re-spectively) were simulated following the procedure in Section. Each dataset was partitioned into training (75%) and testing (25%) sets. Three models were fit using the training set: (1) Model accounting for both clinical predictor and spatial heterogeneity (2) Model accounting for only spatial heterogeneity, and (3) Model accounting for only clinical predictor. The estimated linear predictor 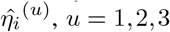 was obtained from the testing set for the *u*th model. At each sample size, mean squared errors *MSE*^(*u*)^ was computed as the average of squared differences between the predicted 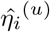 and the “true” linear predictor 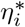such that 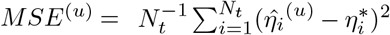, with *N*_*t*_ denoting the number of subjects in the testing set.

We repeated the simulation for 100 iterations and recorded the average MSE for each of the three models across four sample sizes N = 100, 200, 500, 1000 in Table 1. Fig 4 displays the distribution of MSEs for the three models at each noise-added level across the four sample sizes. Note that when there was a low or medium level of subject-specific variation, the separation between the two spatial configurations was clear (Fig. S4 (a)). As a result, by accounting for the impact of such spatial heterogeneity, models (1) and (2) yielded smaller MSEs, as compared to the model (3) across all sample sizes. As the additive noise was greatly increased, the difference in the spatial entropy curves across the two patterns was no longer recognizable. In other words, the spatial heterogeneity was not as predictive as in the previous two scenarios. Consequently, the gain in accounting for spatial impact (i.e., models (1) and (2)) in addition to just clinical predictor (i.e., model (3)) was not as pronounced.

**Table 1.** Mean squared errors (MSE) across three different models, with four sample sizes (N = 100, 200, 500, 1000), and at three levels of additive noise (small, medium, and large). Corresponding standard deviations are recorded in parentheses.

| Noise level | Model | N=100 | N=200 | N=500 | N=1000 |
| --- | --- | --- | --- | --- | --- |
| Small | (1) | 57.09 (17.63) | 60.49 (10.67) | 62.36 (8.25) | 63.80 (6.27) |
|  | (2) | 56.99 (19.81) | 59.76 (13.54) | 63.57 (8.82) | 64.91 (6.48) |
|  | (3) | 137.35 (18.81) | 138.49 (12.80) | 139.09 (8.17) | 139.58 (5.52) |
| Medium | (1) | 84.68 (14.89) | 86.41 (10.63) | 87.25 (6.46) | 87.72 (4.44) |
|  | (2) | 86.20 (15.77) | 87.82 (11.00) | 88.83 (6.85) | 89.27 (4.62) |
|  | (3) | 128.82 (18.65) | 130.79 (12.77) | 131.21 (7.90) | 131.97 (5.46) |
| Large | (1) | 52.76 (15.35) | 57.21 (10.52) | 56.19 (6.66) | 57.79 (4.53) |
|  | (2) | 56.92 (16.58) | 60.85 (11.05) | 59.56 (6.95) | 61.07 (4.67) |
|  | (3) | 70.92(18.77) | 76.53 (13.14) | 74.22 (8.04) | 76.40 (5.69) |

**Fig. 4.**
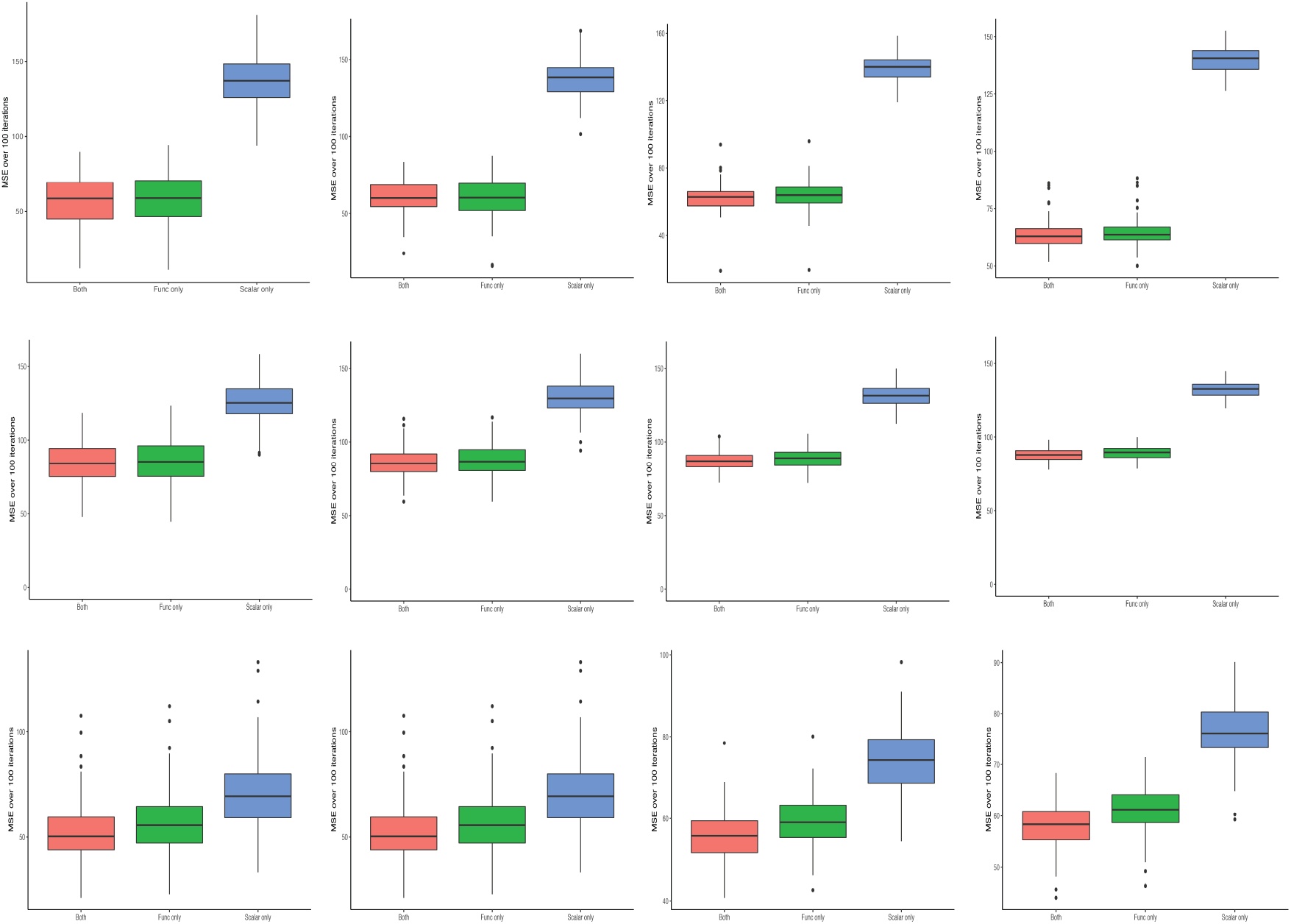
Mean squared errors (MSE) across three different models. Distribution of MSEs for each of the three models: model accounting for both clinical predictor and spatial heterogeneity (red), model accounting for only spatial heterogeneity (green), and model accounting for only clinical predictor (blue). Column from left to right represent four sample sizes: N = 100, 200, 500, and 1000. Rows from top to bottom display three levels of additive noise: small, medium, and large. he lower the MSE, the better the predictive performance.

## Discussion and Conclusions

Spatial architecture of cells in the TME plays a critical role in gaining insights into tumor development, progression, and invasion. Recent advances in imaging technology enable investigators to collect single cell data with an additional dimension of spatial resolution. Conventional metrics typically capture the spatial differential between TME cellular architectures by either exploring the cell distributions in a pairwise fashion or aggregating the heterogeneity across multiple cell distributions without considering for the spatial attribute. As a result, neither has fully accounted for the heterogeneity caused by both cellular diversity and spatial patterns of multiple cell categories. Alternatively, we utilize spatial entropy measures to decompose the conventional Shannon entropy into spatial mutual information and residual entropy, which account for the contribution of space and cellular diversity, respectively, to the overall heterogeneity. Then, we apply functional principal component (FPC) analysis for sparse data to the subject-specific spatial entropy trajectories to estimate the FPC scores. The scores are served as predictors in a Cox regression model to investigate the impact of spatial heterogeneity in the TME on survival outcome, in addition to other clinical variables.

Using the ovarian cancer dataset as a case study, we study the spatial patterns of four different immune cell subsets including CD19+ B cells, CD4+ T cells, CD8+ T cells, and CD68+ macrophages across 114 individuals. After fitting the top three FPC scores into the Cox regression model, we find that the spatial heterogeneity in TME immune compositions has a significant non-zero effect on the overall survival (p = 0.027). The approach is further validated on the TNBC dataset to find the association between the diversity in spatial distributions of tumor cells relative to immune and endothelial cells and risk of mortality. Given a relatively small sample size, such association is not found significant. Additionally, through simulation studies under different spatial configurations, we demonstrate that the proposed method has a higher predictive power by accounting for both clinical effect and the impact of spatial heterogeneity.

In this paper, we utilize the spatial entropy measures to characterize the heterogeneity across distributions of multiple cell types. However, the accuracy of these measures rely heavily on the upstream procedure of cell segmentation and phenotyping. In other words, if cells are not segmented and phenotyped correctly, the estimated spatial entropy values would reflect spurious spatial heterogeneity. One possible solution would be to randomly permutate the cell labels to obtain the empirical distribution of the spatial entropy curves. Then, the mean spatial entropy instead of the observed counterpart would then be used as input for the FPC analysis. The selected FPC scores would be served as scalar predictors in the Cox regression model to investigate the association between spatial heterogeneity of cells and patient overall survival.

## Materials and Methods

### Spatial entropy measures

Let *X* be a random variable denoting a category for each individual cell, with at total of *I* possible categories. Shannon entropy (20) is the expected value of an information function measuring the uncertainty in observing *X* = *x*_*i*_, *i* = 1,…, *I* with the corresponding probability mass function (pmf) as *p*_*X*_ = (*p*(*x*_1_), *p*(*x*_2_), … *p*(*x*_*I*_))^*T*^. The entropy is defined as:

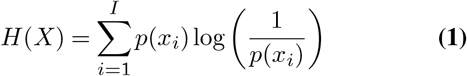

The entropy *H*(*X*) ranges between 0 and log(*I*); and its maximum is achieved when *X* is uniformly distributed. In our context, the maximum entropy is reached when cells of different types are scattered evenly on a given image. Note that the above entropy alone does not account for the role of space. As a result, datasets with identical pmf *p*_*X*_ but different spatial configurations (e.g., strong spatial association vs. complete spatial randomness) yield the same *H*(*X*).

Following (28), we define a new variable *Z* to identify co-occurrences of different pairs of realizations of *X* over space, i.e.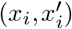, with *i, i*^′^ = 1,… *I*. An assumption is that the order within co-occurrences is neglected. This is reasonable since the interest is to understand the spatial heterogeneity of data over a space, which is usually not assumed to have direction. The number of categories of *Z*, denoted by *R*, where 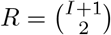. The corresponding pmf is *p* _*X*_ = (*p*(*z*_1_), *p*(*z*_2_), … *p*(*z*_*R*_))^*T*^, where *p*(*z*_*r*_) for *r* = 1,…, *R*, is the probability of observing the *r*th co-occurrence of cells on a given image. In other words, *Z* transforms the information in *X* while introducing a venue for incorporating the idea of spatial neighborhood. The entropy using the newly introduced variable *Z* is defined as:

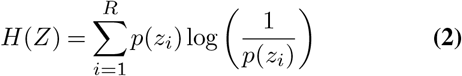

Here, the entropy *H*(*Z*) ranges between 0 and log(*R*). Though the values of *H*(*X*) and *H*(*Z*) might not be exactly the same due to different categories being considered, both capture equivalent information.

In order to properly account for space in an entropy measure, an additional random variable *W* is defined to cover all possible distances at which co-occurrences take place. Denote *w*_*k*_ = (*d*_*k*−1_, *d*_*k*_], with *k* = 1,…, *K* and *d*_*k*_ is a fixed set of distance breaks which are a function of inter-point distances, with *d*_0_ = 0 and *d*_*K*_ being the distance between the two farthest points on a given observation window. These distance breaks *d*_*k*_ can be flexibly chosen depending on the specific applications. The corresponding pmf is denoted by *p*(*W*) = (*p*(*w*_1_), …, *p*(*w*_*K*_))^*T*^, with *p*(*w*_*k*_) being the probability of observing pairs of cells whose corresponding distances fall within the *k*th distance range. At each distance category *w*_*k*_, the probability of observing specific co-occurrences is represented as *p*(*Z* |*w*_*k*_) = (*p*(*z*_1_ |*w*_*k*_), …, *p*(*z*_*R*_ |*w*_*k*_))^*T*^.

By utilizing the two newly defined variables, *Z* and *W*, we are able to decompose the global entropy *H*(*Z*) in Eq. 2 into two components: entropy due space (i.e., spatial information) and the remaining heterogeneity after space has been taken into account (residual entropy), respectively denoted by *SPI*(*Z*) and *H*^*W*^ (*Z*), as follows.

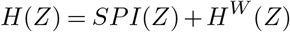

Here, both *SPI*(*Z*) and *H*^*W*^ (*Z*) can be further partitioned by distance range *w*_*k*_, such that

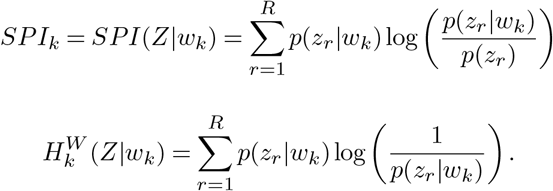

Since our interest lies in investigating the spatial heterogeneity across cellular architectures, we will focus on the spatial entropy *SPI*_*k*_ for the rest of the paper. Fig. 5 displays the distributions of five different cell types including CK+, CD4+ T cells, CD8+ T cells, CD14+, and CD19+ B cells in two representative images (top row) and the corresponding spatial entropy as a function of inter-cell distances (bottom row). Specifically, in the first spatial configuration (top left), the influence of space is rather weak as all cell types seem to scatter evenly across all distance ranges (bottom left). Conversely, the spatial entropy values at short distances are relatively larger due to some small clusters of CD4+ and CD19+ cells.

**Fig. 5.**
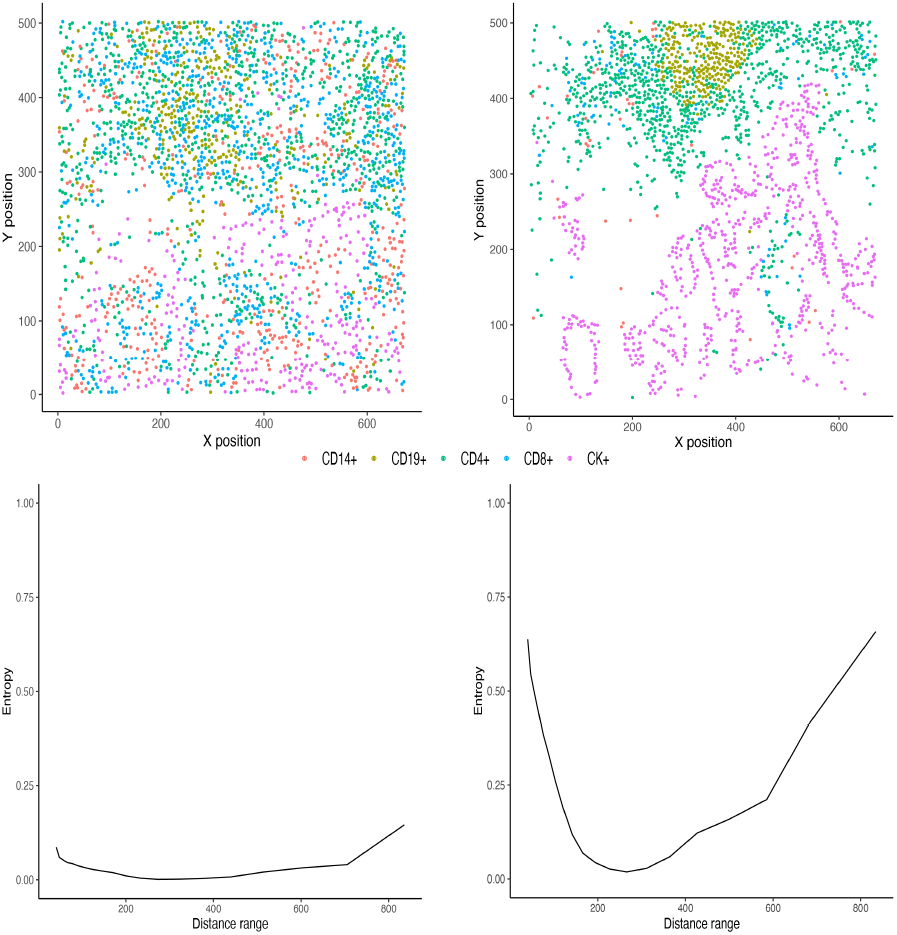
Representative images and corresponding spatial entropy. Top row: two representative images with individual cells of five different types: CD14+, CD19+, CD4+, CD8+, and CK+. Bottom row: corresponding spatial entropy curves at multiple distance ranges.

## Model

### Functional principal component analysis (FPCA) for sparse data

To generalize the entropy measures introduced previously to our context of multiple independent subjects, we suppose that the distance breaks and the associated distance ranges can be independently obtained for each *i*-th subject, denoted as *d*_*ik*_ and *w*_*ik*_ = (*di*(*k*−1), *d*_*ik*_] for *k* = 1,…, *K*_*i*_ and *i* = 1,…, *N*. The spatial entropy for the *i*-th individual are represented as 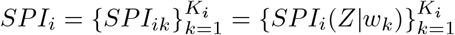 As the number of distance breaks *K*_*i*_ is different across subjects, and typically kept below 20 to ensure the spatial entropy is appropriately computed for each distance range *w*_*ik*_. In other words, if *K*_*i*_ is too large, i.e., *w*_*ik*_ gets close to 0, leading to no point being observed resulting in indefined entropy. Provided the sparse observed entropies for each individual, fitting *SPI*_*i*_ directly into a model as functional covariates as in (31) is not feasible. Therefore, we utilize the approach proposed by Yao et al. (32) to address the sparseness in the observed data through functional principal components (FPC) analysis targeting sparse and irregularly spaced data points.

Assume that each observed spatial entropy *SPI*_*ik*_ for *k* = 1,…, *K*_*i*_, *i* = 1,…, *N* is generated from the underlying, smooth random function *X*(*s*) at a random distance *S*_*ik*_, with known mean function *EX*(*s*) = *µ*(*s*) and covariance function *Cov*(*X*(*s*), *X*(*t*)) = *G*(*s, t*). The domain of *X*(.) is bounded and closed on interval 𝒮. Suppose there is an orthogonal expansion of *G* in terms of eigenfunctions *ϕ*_*l*_ and nonincreasing eigenvalues *λ*_*l*_, such that *G*(*s, t*) = ∑*l* λ_*l*_*ϕ*_*l*_(*s*)*ϕ*_*l*_(*t*) with *t, s* ∈ 𝒮. The *i*-th random curve can be expressed as *X*_*i*_(*s*) = *µ*(*s*) + ∑_*l*_ *ξ*_*il*_*ϕ*_*l*_(*s*), *s* ∈ 𝒮, where *ξ*_*il*_ are uncorrelated random variables with mean 0 and variance 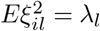 with ∑*l* λ_*l*_ *<* ∞, *λ* ≥ *λ* ≥ …. Then, the observed *SPI*_*ik*_ can be modelled as follows.

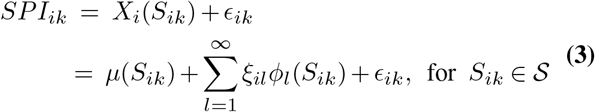

where, *Eϵ*_*ik*_ = 0, *var*(*ϵ*_*ik*_) = *σ*^2^.

Utilizing the local linear smoothers (33), Yao et al. (32) estimate the mean function 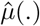 based on pooled data from all subjects. The covariance surface *G*(.) is smoothed using a local quadratic function. The estimated eigenfunctions 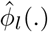 are then obtained by discretizing the smoothed covariance. Details about the smoothing steps can be found in (32). Once the mean function and the eigenfunctions are computed, the FPC scores can be subsequently estimated. Due to the sparseness of the data in which *SPI*_*ik*_ are only observed at discrete random distances *S*_*ik*_, the FPC scores can not be reasonably estimated using traditional numerical integration as 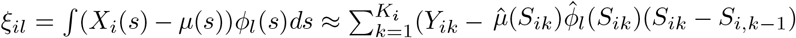, setting *S*_*i*0_ = 0. Alternatively, FPC scores can be estimated through conditional expectation such that

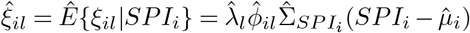

where the (*j, l*)-th element of 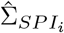 is 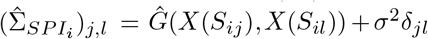 with *δ*_*jl*_ = 1 if *j* = *l*, and 0 otherwise.

### Functional Linear Cox Regression Model

The relationship between the survival distribution and the spatial heterogeneity embedded in the aforementioned FPC scores, in addition to scalar clinical predictors *U*_*i*_ = (*u*_*i*1_, …, *u*_*ip*_)*T* can be investigated using a Cox regression model. Denote *T*_*i*_ and *C*_*i*_ as the survival and censoring times for the *i*-th individual, respectively. Assume that *T*_*i*_ and *C*_*i*_ are independent given *U*_*i*_. Due to right-censoring, we only observe *Y*_*i*_ = *min*(*T*_*i*_, *C*_*i*_), and let *δ*_*i*_ = *I*(*T*_*i*_ ≤ *C*_*i*_) be a censoring indicator. The hazard function for the Cox regression model has the form

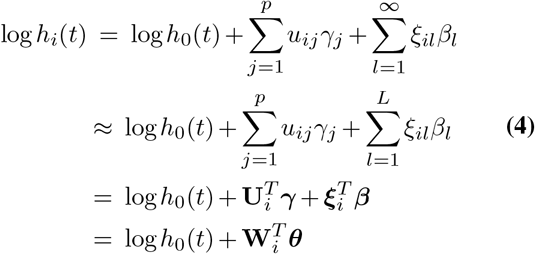

where 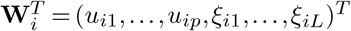, log *h*_*i*_(*t*) is the log hazard at time *t* given scalar covariates *U*_*i*_ and FPC scores *ξ*_*il*_, and log *h*_0_(*t*) is the log baseline hazard function. The truncation number *L* is often chosen such that the resulting FPC scores cumulatively account for at least 95% of observed variance.

The approximate partial log-likelihood function of ***θ***^*T*^ = (***γ***^*T*^, ***β***^*T*^), denoted by *l*(***θ***) is given by

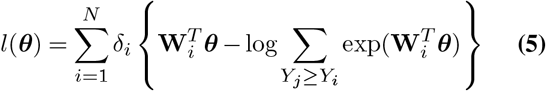

The estimated coefficients 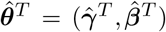 are obtained by maximizing the above partial likelihood using Newton-Raphson procedure.

A likelihood ratio test (LRT) (34) could then be used to investigate the significant association between the spatial information embedded in the FPC scores *ξ*_*il*_ and mortality risk, using the following hypothesis.

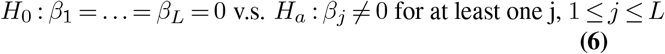

### Data

We used ovarian cancer and triple-negative breast cancer (TNBC) datasets collected using multiplex immunohistochemistry (mIHC) and multiplexed ion beam imaging (MIBI) platforms, respectively, to evaluate the applicability of our proposed model.

### Ovarian cancer data

Tissue microarray (TMA) slides of 132 ovarian cancer patients were stained with antibodies specific for CD3, CD4, CD8, CD19, CD68, cytokeratin, Ki67, pStat, and IER3. The slides were imaged using Vectra 3.0 microscope (Akoya Biosystems) and then segmented and phenotyped using the inForm software. More details can be found in (35). Within the cohort, we excluded 18 patients from the analysis due to missingness of clinical information. Fig. 1 (A) visualizes the distribution of immune cells in the TME including CD19+ B-cells, CD4+ T-cells, CD8+ T-cells, and CD68+ macrophages in four representative images.

### Triple-negative breast cancer data

TNBC biopsies were compiled into a tissue microarray (TMA) slides and stained with 36 antibodies targeting regulators of immune activation such as PD1, PD-L1, etc. The slides were imaged using the multiplexed ion beam imaging (MIBI) mass spectrometer. Details about nuclear segmentation and cell phenotyping of the 41 images can be found in (8). Within this cohort, two patients did not have clinical information available regarding survival outcomes, and one patient’s imaging data was corrupted with a high level of noise. As a result, only data of 38 patients were considered in the analysis.

## ACKNOWLEDGEMENTS

We thank the Human Immune Monitoring Shared Resource and support of the University of Colorado Human Immunology and Immunotherapy Initiative for their expert assistance in multiplex IHC and generation of the ovarian dataset.

## Supplementary Information

**Fig. S1:**
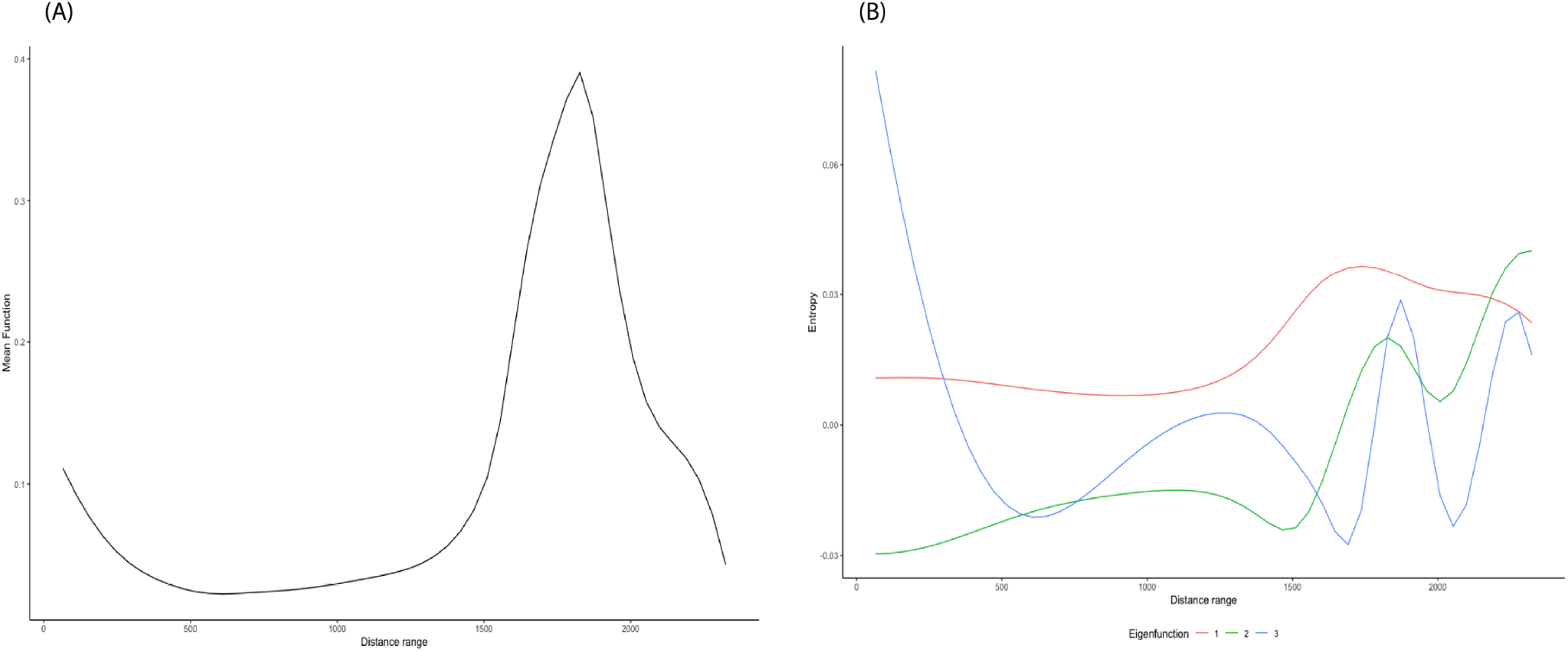
Mean function and eigenfunctions from FPC analysis on SPI curves in ovarian cancer dataset.

**Fig. S2:**
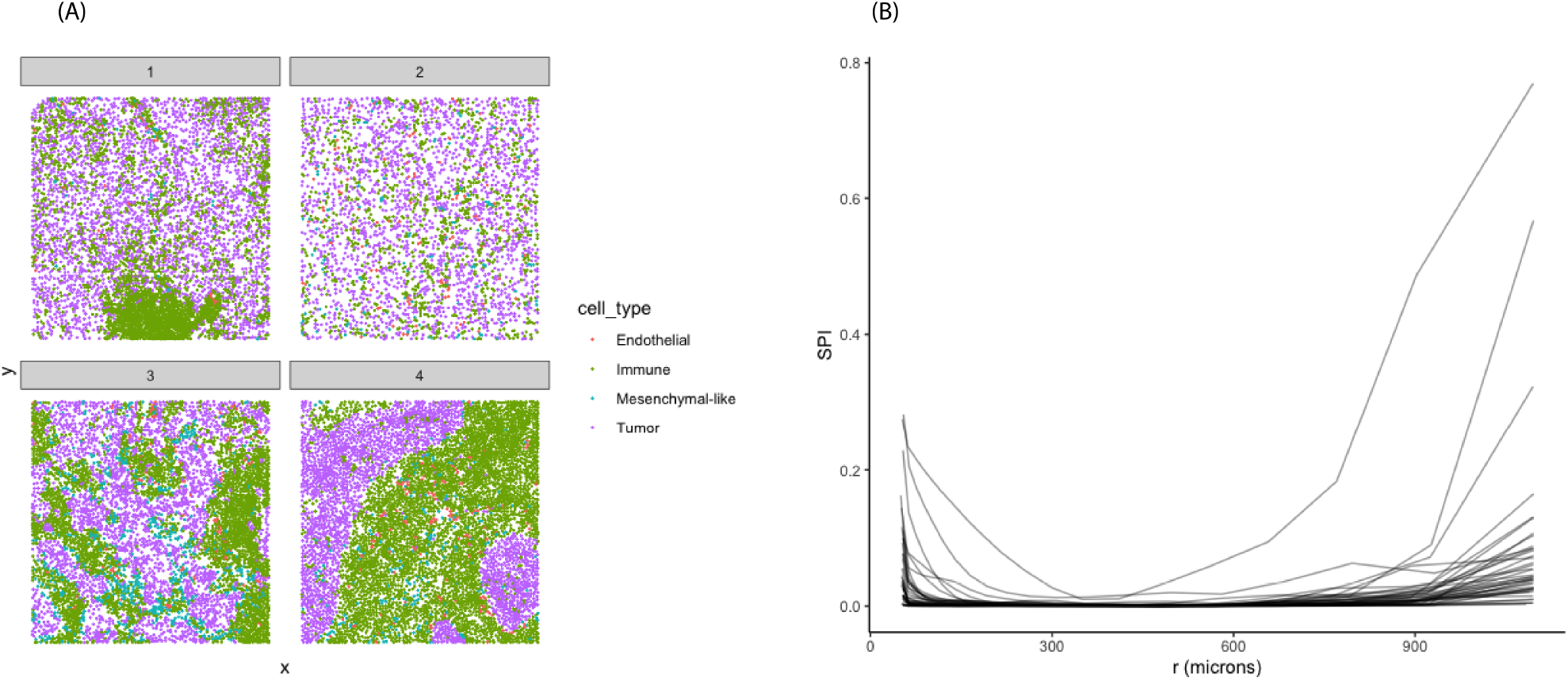
Representative images and corresponding spatial entropy measures across four cell types: endothelial, immune, mesenchymal-like, and tumor. High SPI values indicate clustering patterns while small values occur when cell of different types are scattered more evenly.

**Fig. S3:**
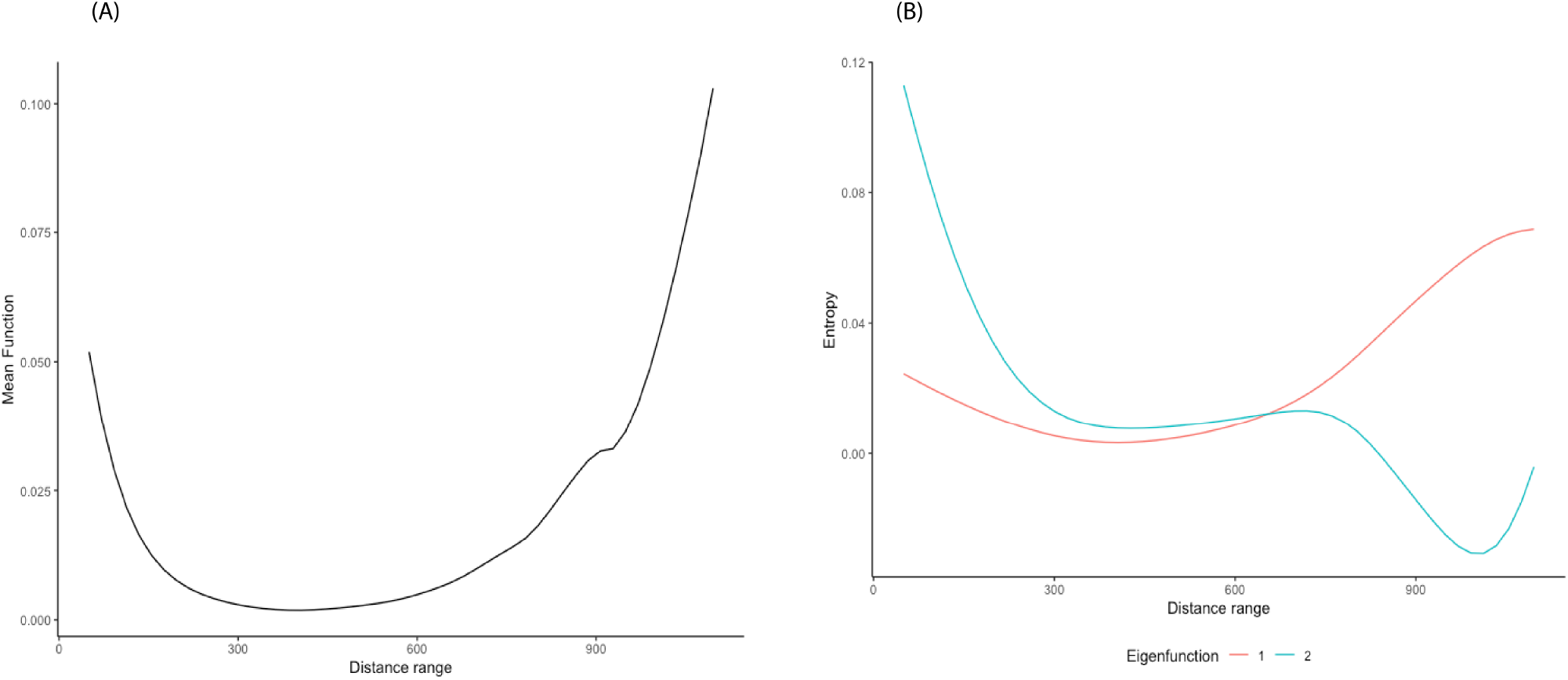
Mean function and eigenfunctions from FPC analysis on SPI curves in TNBC dataset

**Fig. S4:**
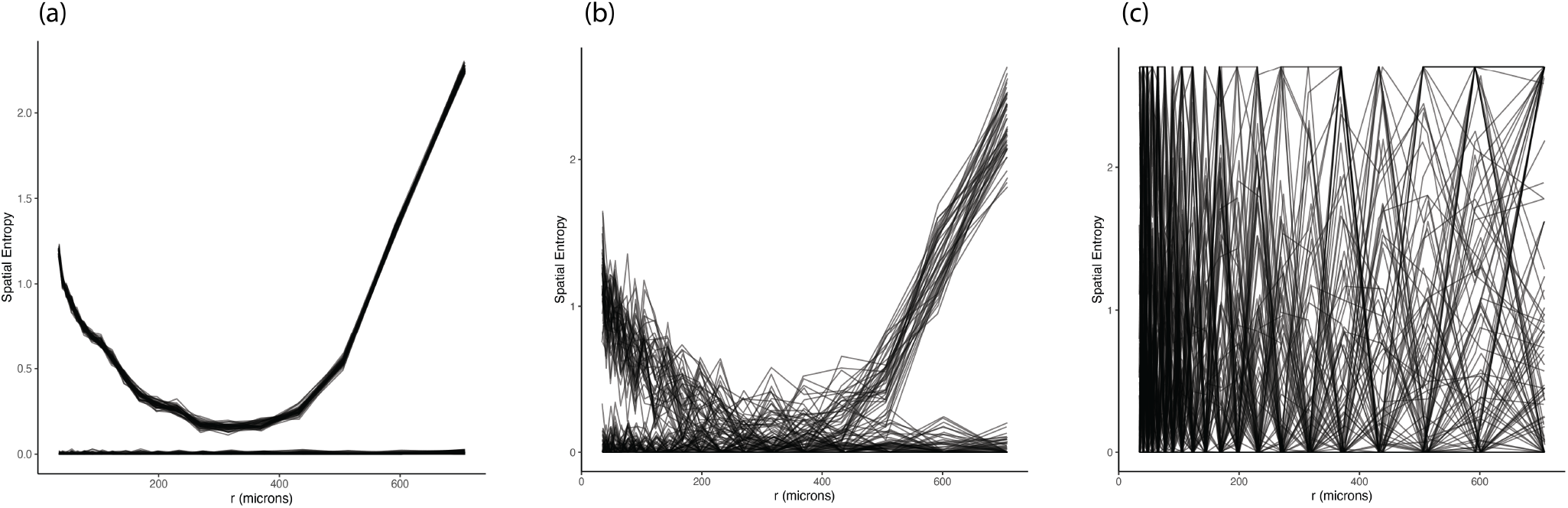
Simulation scenarios: (a) low additive noise. (b) medium additive noise. (c) large additive noise. Three levels of noise were added to the reference SPI curves (clustered vs. random) to generate subject-specific SPI curves.

## Notes

### Competing Interest Statement

The authors have declared no competing interest.

