## Supplementary Information for "FunSpace: A functional and spatial analytic approach to cell imaging data using entropy measures"

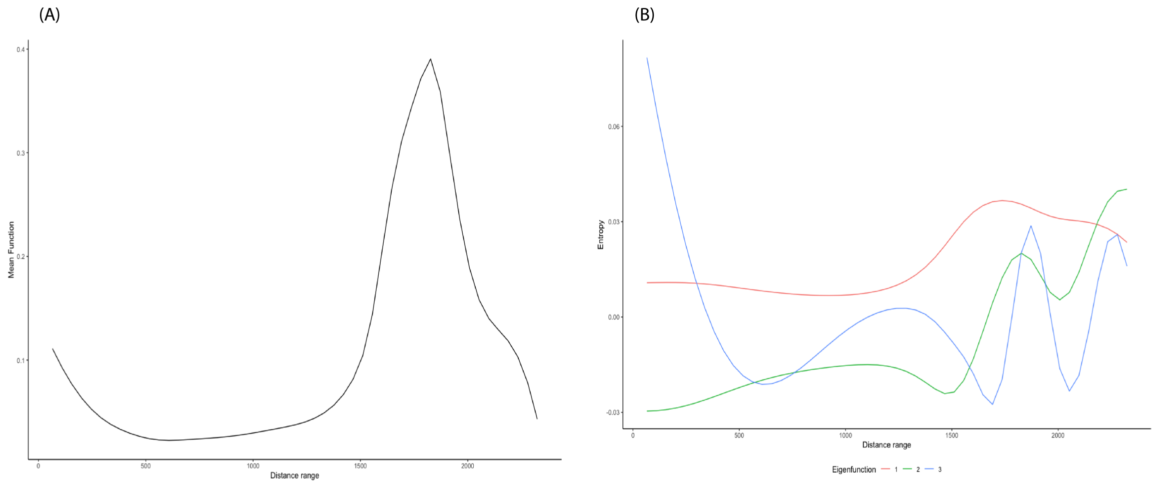


Fig. S 1: Mean function and eigenfunctions from FPC analysis on SPI curves in ovarian cancer dataset.

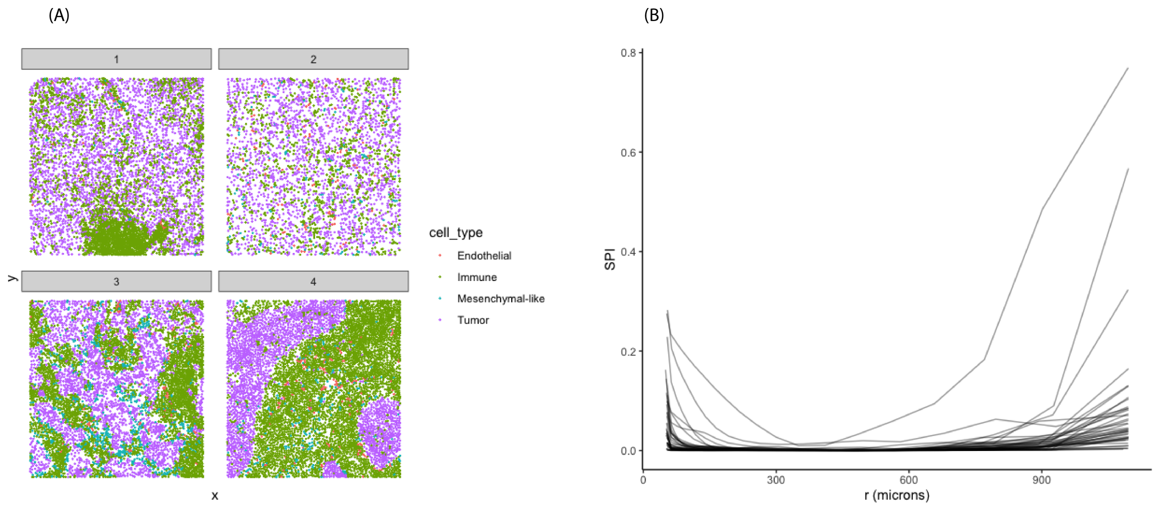


Fig. S 2: Representative images and corresponding spatial entropy measures across four cell types: endothelial, immune, mesenchymal-like, and tumor. High SPI values indicate clustering patterns while small values occur when cell of different types are scattered more evenly.

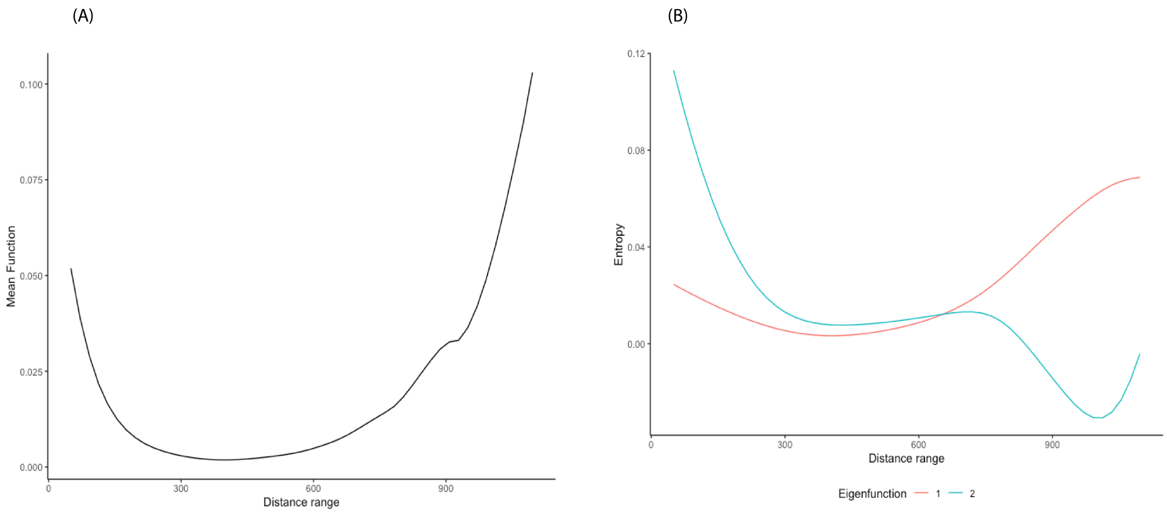


Fig. S 3: Mean function and eigenfunctions from FPC analysis on SPI curves in TNBC dataset

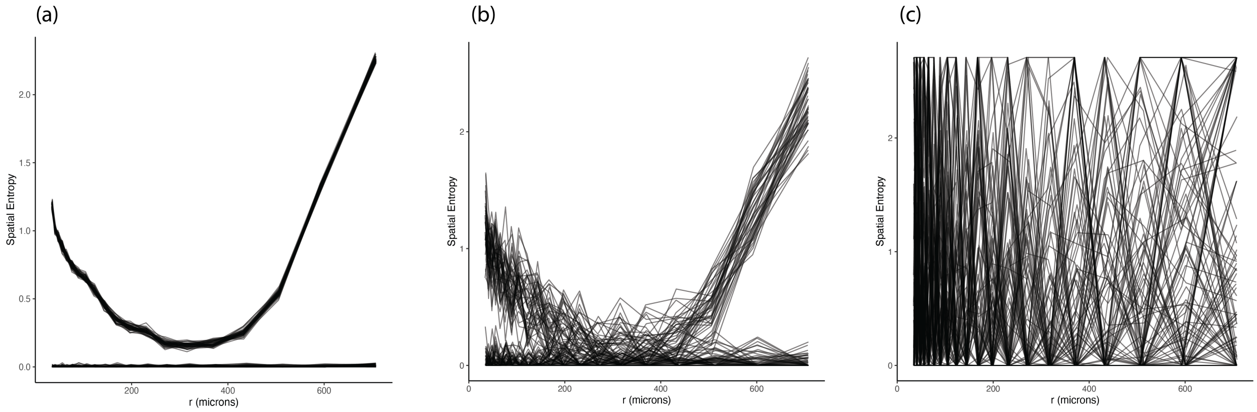


Fig. S 4: Simulation scenarios: (a) low additive noise. (b) medium additive noise. (c) large additive noise. Three levels of noise were added to the reference SPI curves (clustered vs. random) to generate subject-specific SPI curves.
